# Physiological role of the 3’IgH CBEs super-anchor in antibody class switching

**DOI:** 10.1101/2020.11.25.398404

**Authors:** Xuefei Zhang, Hye Suk Yoon, Aimee M. Chapdelaine-Williams, Nia Kyritsis, Frederick W. Alt

**Author notes:** Corresponding Author: Frederick W. Alt, Howard Hughes Medical Institute, Boston Children’s Hospital, Program in Cellular and Molecular Medicine, Department of Genetics, Harvard Medical School, 300 Longwood Avenue, Boston, MA 02115, USA.

## Abstract

IgH class switch recombination (CSR) replaces Cμ constant region (C_H_) exons with one of six downstream C_HS_ by joining transcription-targeted DSBs in the Cμ switch (S) region to DSBs in a downstream S region. Chromatin loop extrusion underlies fundamental CSR mechanisms including 3’IgH regulatory region (3’IgHRR)-mediated S region transcription, CSR center formation, and deletional CSR joining. There are ten consecutive CTCF binding elements (CBEs) downstream of the 3’IgHRR, termed the “3’IgH CBEs”. Prior studies showed that deletion of eight 3’IgH CBEs did not detectably affect CSR. Here, we report that deletion of all 3’IgH CBEs impacts, to varying degrees, germline transcription and CSR of upstream S regions, except Sγ1. Moreover, deletion of all 3’IgH CBEs rendered the 6kb region just downstream highly transcribed and caused sequences within to be aligned with Sμ, broken, and joined to form aberrant CSR rearrangements. These findings implicate the 3’IgH CBEs as a critical insulator for focusing loop extrusion-mediated 3’IgHRR transcriptional and CSR activities on upstream C_H_ locus targets.

**Significance:** B lymphocytes change antibody heavy chain (IgH) isotypes by a recombination/deletion process called IgH class switch recombination (CSR). CSR involves introduction of DNA breaks into a donor switch (S) region and also into one of six downstream S regions, with joining of the breaks changing antibody isotype. A chromatin super-anchor, of unknown function, is located just downstream of the IgH locus. We show that complete deletion of this super-anchor variably decreases CSR to most S regions and creates an ectopic S region downstream of IgH locus that undergoes aberrant CSR-driven chromosomal rearrangements. Based on these and other findings, we conclude that the super-anchor downstream of IgH is a critical insulator for focusing potentially dangerous CSR rearrangements to the IgH locus.

## Introduction

Mature B cells undergo immunoglobulin (Ig) heavy chain (IgH) class switch recombination (CSR) to change the constant region of IgH chains and modulate antibody effector functions (1, 2). Transcription from IgH V(D)J exons runs through proximal Cμ exons that specify IgM antibodies. Upon activation, B cells undergo CSR to replace Cμ with one of six sets of constant region exons (C_Hs_) to produce other antibody isotypes (IgG, IgE or IgA) (2, 3). The intronic enhancer (iEμ), located at the 5’ end, and 3’IgH regulatory region (3’IgHRR) super-enhancer, located at the 3’ end, of the IgH constant region locus both contribute to CSR (4–9). The 3’IgHRR plays critically important roles in CSR by interacting with I-promoters upstream of targeted S regions to activate C_H_ transcription in a cytokine and activation specific manner (10, 11). Then, transcriptionally-targeted activation-induced cytidine deaminase (AID) (12) initiates DSBs in downstream acceptor S regions that can join in deletional orientation to AID initiated DSBs in the donor S upstream of Cμ (Sμ) (1). Insertion of active promoters in various C_H_ locus sites inhibits I-promoter activation and CSR to upstream promoters (except Iγ1) but not downstream promoters, suggesting that linear competition over 100kb distances of I-promoters for 3’IgHRR activation contributes to CSR regulation (13).

Regulated chromatin loop extrusion provides mechanistic underpinnings for the overall CSR mechanism by promoting synapsis of enhancers, promoters, S regions and DSB ends necessary for productive, deletional CSR (11). In naïve B cells, cohesin is loaded onto the chromatin around the IgH enhancers (iEμ or 3’IgHRR) and then extrudes the 3’IgHRR into proximity with iEμ-Sμ region to form a dynamic CSR center (CSRC) containing donor Sμ and the two involved enhancer regions (11). Besides loading cohesin, these enhancer regions appear to also function as dynamic loop extrusion impediments that contribute to formation of a CSRC (11). In CSR-activated B cells, loop extrusion also brings cytokine/activator primed I promoters into the CSRC where they can be further transcriptionally activated by IgH enhancers, resulting in further cohesin loading, loop extrusion, and alignment the transcribed acceptor S region with donor Sμ region for S-S synapsis and AID targeting (11). For the next joining step, it has been implicated that cohesin rings put tension on the S regions synapsed in the CSRC to promote AID-initiated DSBs ends in donor and acceptor S regions to be dominantly joined in deletional orientation for CSR. Based on this model, after AID initiates DSBs, one or both break ends within an S region will be reeled into an opposing cohesin ring where extrusion process is stalled. Then a break on the other S region will have the same fate, eventually aligning the break-ends for deletional CSR joining (11). The model also has been proposed to explain how DSBs within ectopic S regions (non-S region sequences) formed by CBE insertions into the CH region also are synapsed with Sμ in the CSRC, after which their infrequent AID-initiated DSBs are joined in deletional orientation to AID-initiated Sμ DSBs (11).

The ten consecutive CTCF binding elements (CBEs) downstream of 3’IgHRR, variously termed the 3’IgH CBEs or the 3’IgH locus super-anchor have been speculated to function as an insulator at 3’end of IgH locus (14–16). However, deletion of the first eight of the ten 3’IgH CBEs showed little effect on CSR in mice (17). Deletion of all ten 3’IgH CBEs in CH12F3 cells revealed modest effects on the transcription and CSR to Cα (15). However, transcription of the 3’IgHRR and the Cα region appears to be constitutively activated in CH12F3 cells versus normal B cells, being much more robust and extending more than 20kb downstream through the 3’IgH CBEs and beyond (11). Thus CH12F3 cells do not necessarily provide an accurate model for studying potential roles of 3’IgH CBEs in physiological CSR. As complete deletion of the 3’IgH CBEs in normal B cell CSR had not yet been assayed, it has remained unknown as to whether or not the 3’IgH CBEs play any potential direct or indirect roles in the physiological CSR process. Here, we describe experiments in which all ten 3’IgH CBEs were deleted (“complete 3’IgH CBEs-deleted”) on both alleles in ES cells that were then used for RAG2-deficient blastocyst complementation (RDBC) (18) to generate chimeric mice in which all mature B cells harbor the complete 3’IgH CBEs-deletion. Our current studies of CSR in complete 3’IgH CBEs-deleted B cells indeed revealed roles for the 3’IgH CBEs in physiological CSR and clear-cut function for 3’IgH CBEs as a physiological CSR insulator.

## Results

### Complete 3’IgH CBEs deletion decreases CSR to most S regions

While previous work indicated that deletion of eight of the ten CTCF binding sites of the 3’IgH CBEs had little effect on class switching (17), it remained possible that the remaining two 3’IgH CBEs might suffice for mediating potential CSR functions. Therefore, to assess potential physiological roles of 3’IgH CBEs in CSR, we deleted all ten of the 3’IgH CBEs in ES cells (Fig. 1A; Fig. S1 A and B) and used our RDBC system (18) to generate chimeric mice in which all mature B cells derive from the donor 3’IgH CBEs-deleted ES cells. We isolated the primary splenic B cells from the WT and 3’IgH CBEs-deleted RDBC chimeras and stimulated the cells for 96 hours with either αCD40/IL4 to induce class-switching to Sγ1 and Sε, or with LPS/αIgD-dextran to induce CSR to Sγ3, Sγ2b, and Sγ2a (19). Subsequently, we assayed for CSR by CSR-HTGTS-seq (11, 19) (Fig. S1C). Approximately 75% of splenic B cells activated with αCD40/IL4 switched to Sγ1 and approximately 10% switched to Sε (Fig. 1 B and C; Fig. S1D). While the complete 3’IgH CBEs deletion had no significant effect on CSR to Sγ1, it decreased CSR to Sε to about 30% of normal levels (from 10.4 to 2.8%) (Fig. 1 B and C; Fig. S1D; Table S1). In LPS/αIgD-dextran treated splenic B cells, the complete 3’IgH CBEs deletion modestly decreased CSR to Sγ3 to about 60% of WT B cell levels (from 14.1% to 8.1%) and CSR to Sγ2b to about 75% of WT B cell levels (from 21.3% to 16.3%) (Fig. 2 A and B; Fig. S1E; Table S2). However, CSR to Sγ2a was substantially decreased to about 15% (from 22.9% to 3.4%) of WT B cell levels (Fig. 2 A and B; Fig. S1E; Table S2).

**Figure 1.**
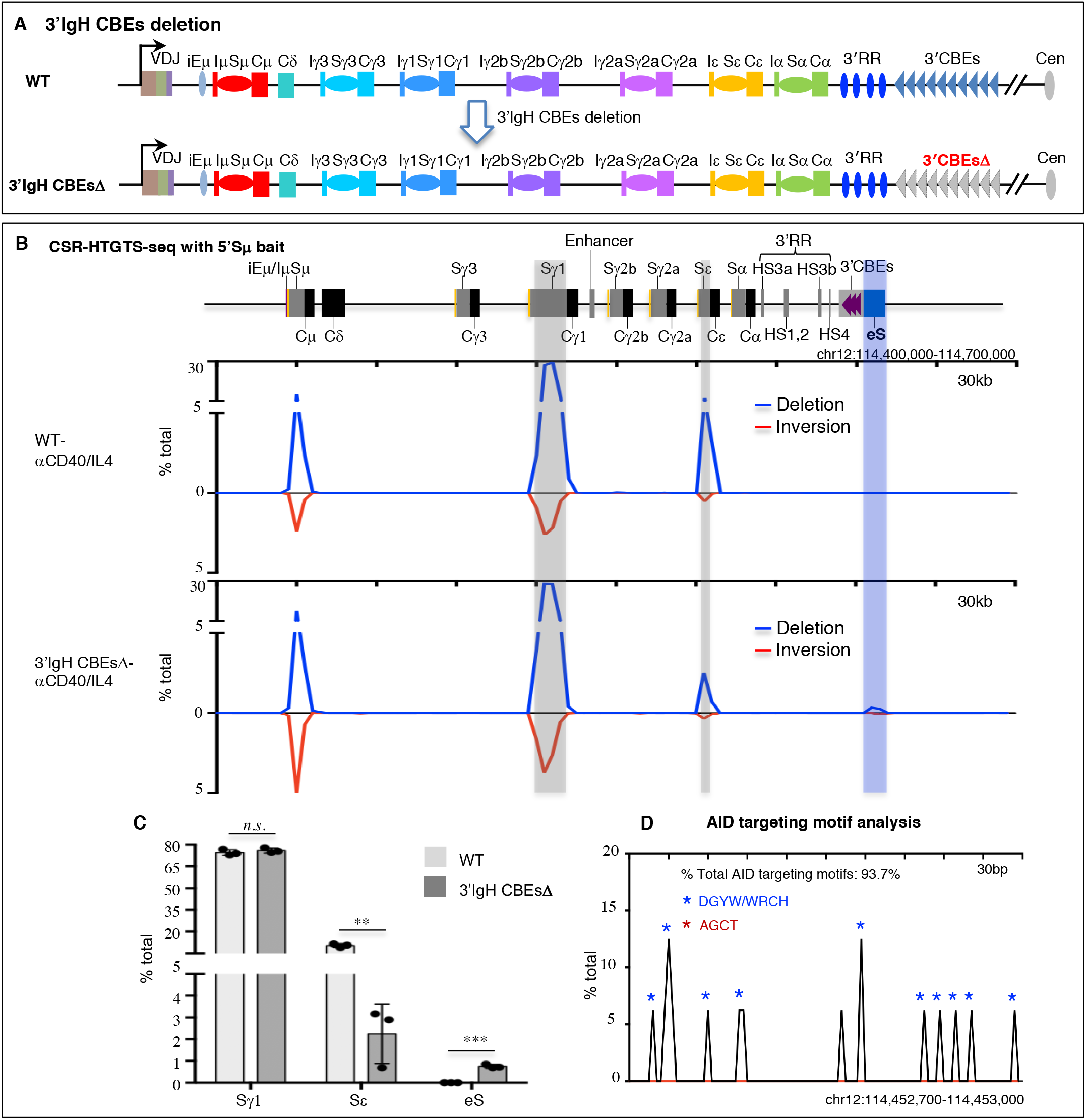
3’IgH CBEs deletion decreases Sε CSR and creates an eS region for aberrant translocation in αCD40/IL4-stimulated B cells. (A) Schematic of IgH locus from iEμ to 3’IgH CBEs and illustration the generation of 3’IgH CBE-deleted ES cells. (B) CSR-HTGTS-seq analysis of break joining between 5’Sμ and downstream acceptor S or non-S regions in WT and 3’IgH CBEs-deleted splenic B cells stimulated with αCD40/IL4. The blue line indicates deletional joining and the red line indicates inversional joining. Grey bars highlight the Sγ1 and Sε. A blue bar highlight the ectopic S region (labeled as “eS”) just downstream of 3’IgH CBEs. (C) Bar graph shows percentages of junctions located in different S regions or eS region from WT and 3’IgH CBEs-deleted splenic B cells stimulated with αCD40/IL4. Data represents mean ± s.d. from three independent repeats. *P* values were calculated via an unpaired two-tailed *t*-test, *n.s*. indicates *p*> 0.05, ** p≤ 0.01, *** p≤ 0.001. The raw data for this bar graph is summarized in Table S1. (D) AID-targeting-motif analysis for the junctions located in a 300bp region within AID-targeted eS region from 3’IgH CBEs-deleted splenic B cells stimulated with αCD40/IL4. Blue asterisks indicate DGYW/WRCH motifs and red asterisks indicate AGCT motifs.

**Figure 2.**
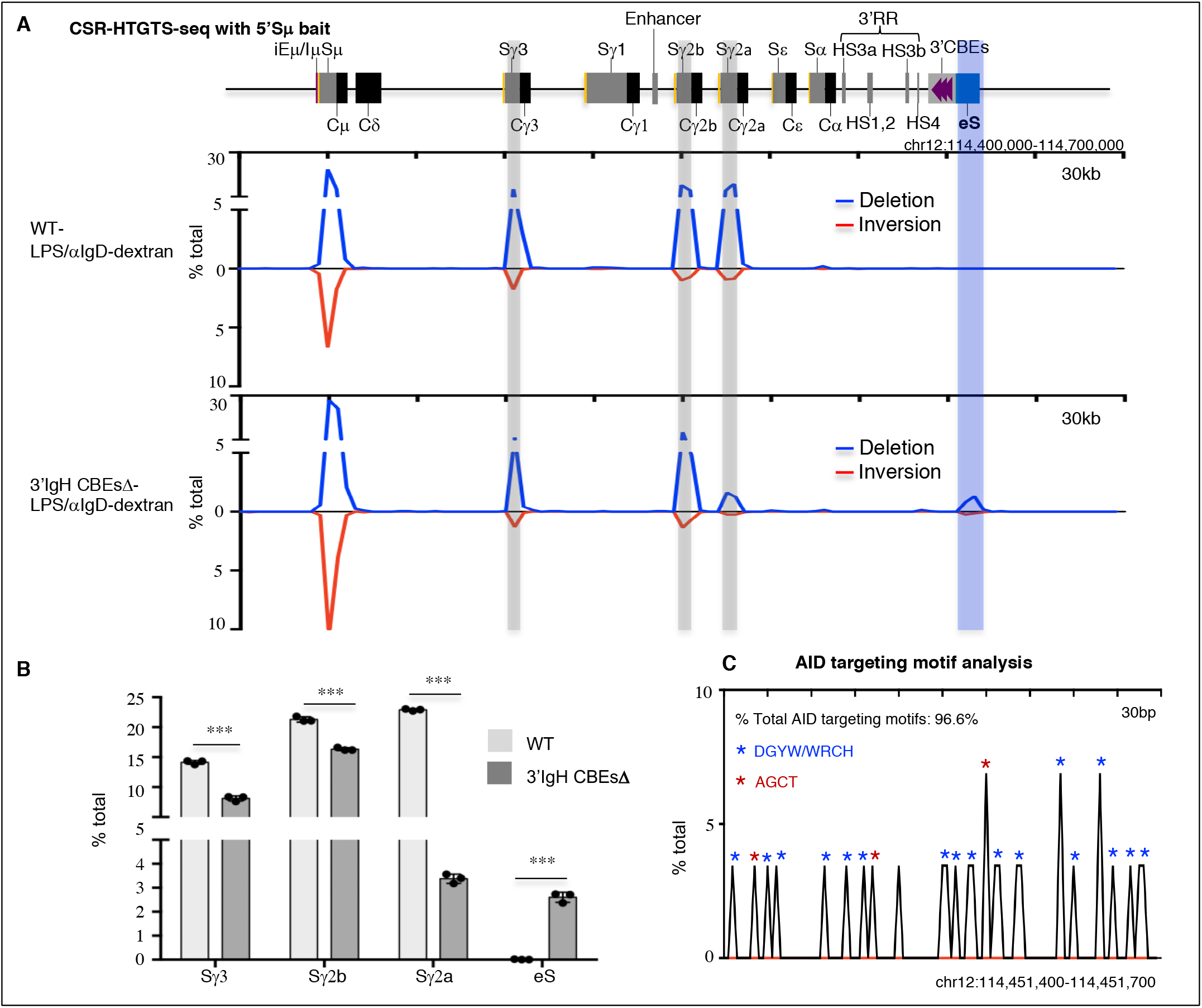
3’IgH CBEs deletion decreases Sγ3, Sγ2b and Sγ2a CSR and creates an eS region for aberrant translocation in LPS/αIgD-dextran-stimulated B cells. (A) CSR-HTGTS-seq analysis of break joining between 5’Sμ and downstream acceptor S or non-S regions in WT and 3’IgH CBEs-deleted splenic B cells stimulated with LPS/αIgD-dextran. The blue line indicates deletional joining and the red line indicates inversional joining. Grey bars highlight the Sγ1, Sγ2b and Sγ2a. A blue bar highlight the ectopic S region (labeled as “eS”) just downstream of 3’IgH CBEs. (B) Bar graph shows percentages of junctions located in different regions from WT and 3’IgH CBEs-deleted splenic B cells stimulated with LPS/αIgD-dextran. Data represents mean ± s.d. from three independent repeats. *P* values were calculated via an unpaired two-tailed *t*-test, *** p≤ 0.001. The raw data for this bar graph is summarized in Table S2. (C) AID-targeting-motif analysis for the junctions located in a 300bp region within AID-targeted eS region from 3’IgH CBEs-deleted splenic B cells stimulated with LPS/αIgD-dextran. Blue asterisks indicate DGYW/WRCH motifs and red asterisks indicate AGCT motifs.

### The complete 3’IgH CBEs deletion decreases the transcription of most S regions

CSR is targeted to particular downstream acceptor S regions by activation/cytokine-induced transcription through them from an I region promoter that lies just upstream of each of them. Induction of transcription from all I region promoters, except that of Iγ1 (8, 13, 20, 21), is dependent on interactions with the 3’IgHRR enhancers. We used GRO-seq to assess the transcription across the C_H_-containing portion of the IgH locus and immediately downstream sequences in WT and 3’IgH CBEs-deleted splenic B cells with or without CSR activation for 96 hours. To obviate effects of CSR events on transcription patterns, we also deleted AID in both the WT and 3’IgH CBEs-deleted ES cells before use for RDBC. Treatment with αCD40/IL4, as expected (11, 19, 22), induced WT B cells to transcribe across Iγ1-Sγ1 and Iε-Sε (Fig. 3A; Fig. S2). In approximate correspondence to CSR effects, the 3’IgH CBEs deletion had no significant effects on Iγ1-Sγ1 transcription but reduced Iε-Sε transcription to about 40% of WT B cell levels (Fig. 3 A and B; Fig. S2A; Table S1). Activation of splenic B cells with LPS/αIgD-dextran, again as expected (19), induced transcription across Iγ3-Cγ3, Iγ2b-Cγ2b and Iγ2a-Cγ2a (Fig. 3C; Fig. S3). Compared to WT B cells levels for this treatment, the 3’IgH CBEs deletion decreased Iγ3-Cγ3 transcription to about 15%, Iγ2b-Cγ2b transcription to about 30%, and Iγ2a-Cγ2a transcription to about 13% (Fig. 3 C and D; Fig. S3; Table S2). While the reduction in transcription levels across the Iγ3-Cγ3, Iγ2b-Cγ2b, and Iγ2a-Cγ2a again do not absolutely reflect the reduction in CSR levels to these C_H_ units in the LPS/αIgD-dextran treated WT and 3’IgH CBEs-deleted B cells, the general trends are similar. In this regard, we do not know the threshold of transcription for each S region required to promote given levels of CSR, which also is influenced by S region sequence composition or length, among other potential factors (23, 24). Thus, we do not necessarily expect precise correspondence between CSR and transcription levels of a particular S region.

**Figure 3.**
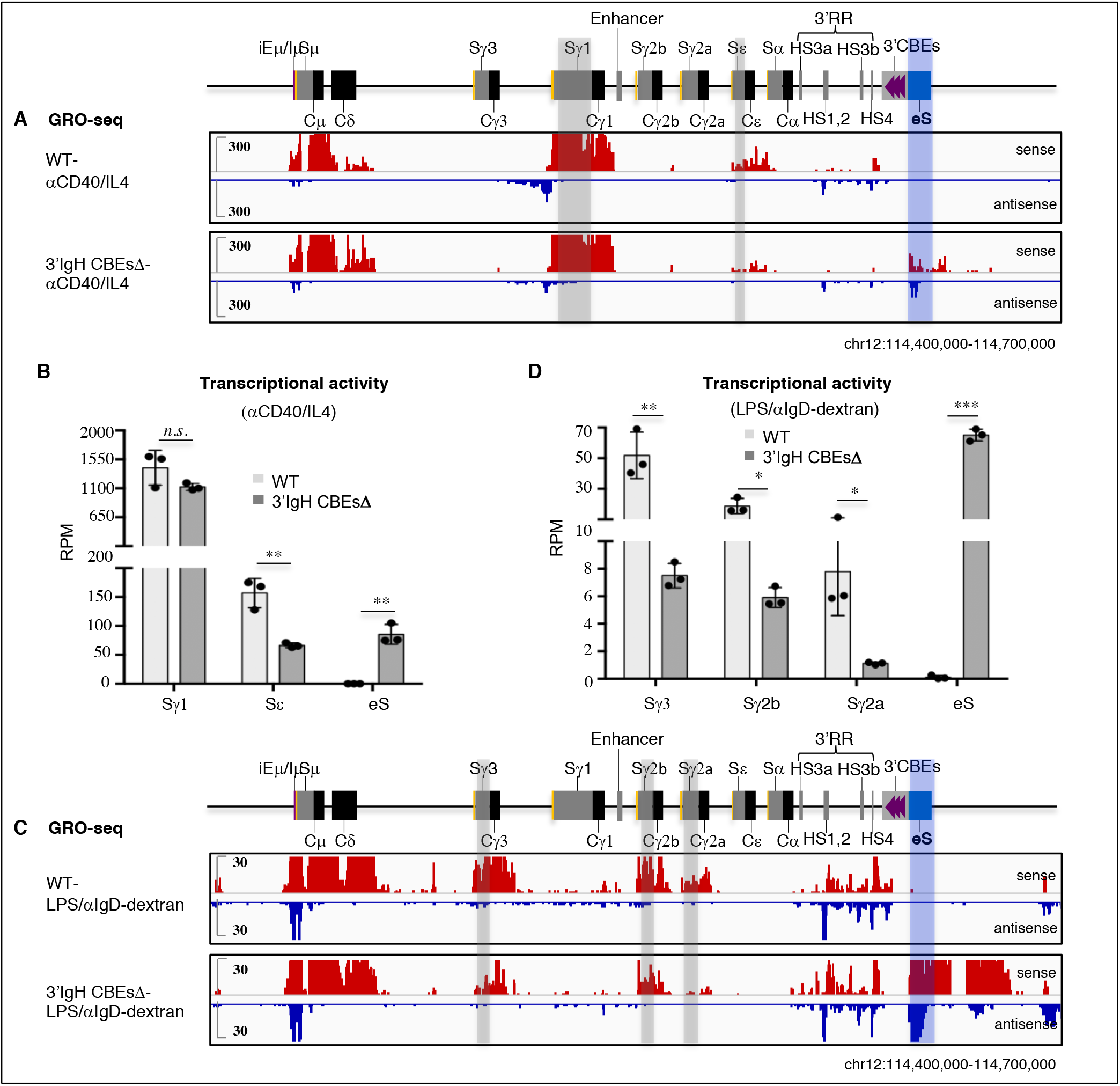
3’IgH CBEs deletion decreases transcription of the most upstream S region transcription and induces the transcription of the eS region. (A) GRO-seq profiles of IgH locus from WT and 3’IgH CBEs-deleted splenic B cells stimulated with αCD40/IL4. Sense transcription is shown above in red and antisense transcription is shown below in blue lines. Grey bars highlight the Sγ1 and Sε. A blue bar highlight the ectopic S region (labeled as “eS”) just downstream of 3’IgH CBEs. (B) Bar graph shows GRO-seq transcriptional activity (calculated as RPM) of the different indicated S regions and eS region in αCD40/IL4 stimulated WT and 3’IgH CBEs-deleted splenic B cells. Data represents mean ± s.d. from three independent repeats. *P* values were calculated via unpaired two-tailed *t*-test, *n.s*. indicates *p*> 0.05, ** p≤ 0.01. The raw data for this bar graph is summarized in Table S1. (C) GRO-seq profiles of IgH locus from WT and 3’IgH CBEs-deleted splenic B cells stimulated with LPS/αIgD-dextran. Sense transcription is shown above in red and antisense transcription is shown below in blue lines. Grey bars highlight the Sγ1, Sγ2b and Sγ2a. A blue bar highlights the ectopic S region (labeled as “eS”) just downstream of 3’IgH CBEs. (D) Bar graph shows GRO-seq transcriptional activity (calculated as RPM) of the different indicated S regions and eS region in LPS/αIgD-dextran stimulated WT and 3’IgH CBEs-deleted splenic B cells. Data represents mean ± s.d. from three independent repeats. *P* values were calculated via unpaired two-tailed *t*-test, *n.s*. indicates *p*> 0.05, * p≤ 0.05, ** p≤ 0.01, *** p≤ 0.001. The raw data for this bar graph is summarized in Table S2.

### The complete 3’IgH CBEs deletion decreases S region synapsis with the CSRC

Our prior studies, which employed the highly sensitive 3C-HTGTS chromatin interaction method (11) indicated that the transcribed iEμ-Sμ and the 3’IgHRR regions serve as a dynamic loop extrusion impediments that can promote CSRC formation and S-S synapsis in CSRC to promote CSR. To assess potential effects of the complete 3’IgH CBEs deletion on S region synapsis in the CSRC, we perform 3C-HTGTS with and iEμ-Sμ bait in activated WT and 3’IgH CBEs-deleted splenic B cells. As noted previously, portions of Sμ and certain other S regions cannot be mapped by this assay due to lack of requisite NlaIII restriction endonuclease sites; and, thus, their interactions must be inferred from mappable sequences within them (11). In αCD40/IL4-stimulated B cells, the iEμ-Sμ locale significantly interacts with Sγ1 and Sε locales (Fig. 4A; Fig. S4A). In this regard, the 3’IgH CBEs complete deletion had no significant effects on Sμ-Sγ1 synapsis; while it modestly reduced Sμ-Sε synapsis to about 75% of WT B cell levels (Fig. 4A; Fig. S4A; Table S1). In LPS/αIgD-dextran-stimulated B cells, iEμ-Sμ locale had relatively less interaction with the Sγ3, Sγ2b and Sγ2a (Fig. 4E; Fig. S4B); however the 3’IgH CBEs deletion significantly decreased Sμ-Sγ3 synapsis to about 45%, Sμ-Sγ3 synapsis to about 60%, and Sμ-Sγ3 synapsis to about 65% of WT B cell levels (Fig. 4E; Fig. S4B; Table S2). Again, the trend of these reductions is in the same direction as CSR, but also is subject, beyond mapping issues with some core S region sequences to the same comparison issues mentioned for correlation of S region transcription levels for CSR levels above.

**Figure 4.**
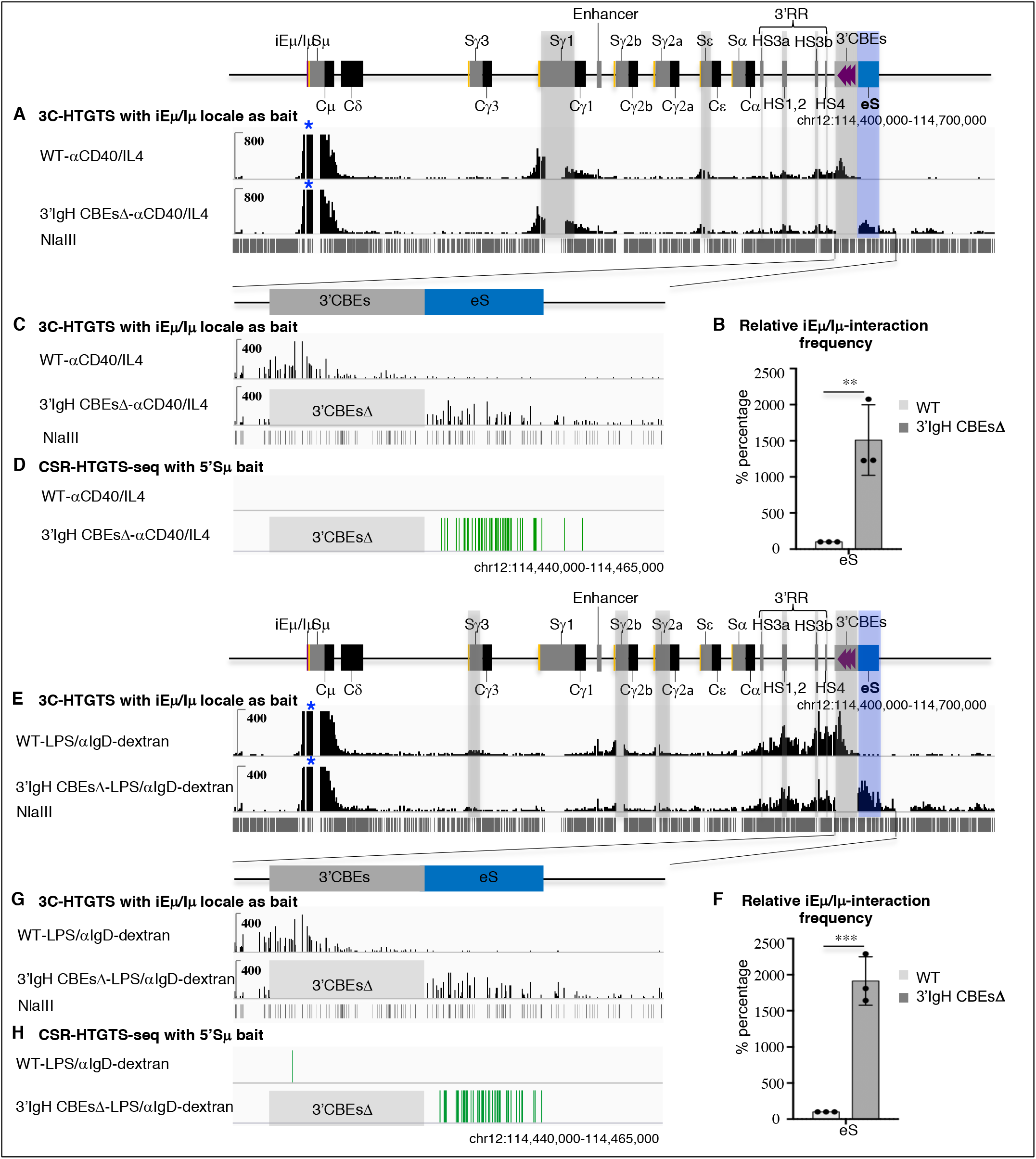
3’IgH CBEs deletion decreases most S-S synapsis and induces Sμ-eS synapsis for aberrant rearrangement. (A) 3C-HTGT analysis of αCD40/IL4-stimulated WT and 3’IgH CBEs-deleted splenic B cells using iEμ/Iμ locale as bait (blue asterisk). Grey bars highlight the Sγ1, Sε, HS3a, HS1,2, HS3b, HS4 and 3’IgH CBEs. Blue bar highlight the ectopic S region (labeled as “eS”) just downstream of 3’IgH CBEs. (B) Bar graph shows the relative iEμ-Sμ interaction frequency with eS in αCD40/IL4-stimulated splenic B cells. Data represents mean ± s.d. from three independent repeats. *P* values were calculated via unpaired two-tailed *t*-test, ** p≤ 0.01. The raw data for this bar graph is summarized in Table S1. (C) Magnified 3C-HTGTS profiles in (A) to better reveal interaction patterns for the 3’IgH CBEs and eS region in αCD40/IL4-stimulated splenic B cells. (D) CSR-HTGTS-seq with 5’Sμ bait to show the rearrangement within the eS region from αCD40/IL4-stimulated WT and 3’IgH CBEs-deleted splenic B cells. (E) 3C-HTGTS analysis of LPS/αIgD-dextran-stimulated WT and 3’IgH CBEs-deleted splenic B cells using iEμ/Iμ locale as bait (blue asterisk). Grey bars highlight the Sγ3, Sγ2b, Sγ2a, HS3a, HS1,2, HS3b, HS4 and 3’IgH CBEs. Blue bar highlight the ectopic S region (labeled as “eS”) just downstream of 3’IgH CBEs. (F) Bar graph shows the relative iEμ-Sμ interaction frequency with eS in LPS/αIgD-dextran-stimulated splenic B cells. Data represents mean ± s.d. from three independent repeats. *P* values were calculated via unpaired two-tailed *t*-test, *** p≤ 0.001. The raw data for this bar graph is summarized in Table S2. (G) Magnified 3C-HTGTS profiles in (E) to better reveal the interaction patterns for the 3’IgH CBEs and eS region in LPS/αIgD-dextran-stimulated splenic B cells. (H) CSR-HTGTS-seq with 5’Sμ bait to show the rearrangement within the eS region from LPS/αIgD-dextran-stimulated WT and 3’IgH CBEs-deleted splenic B cells.

### 3’IgH CBEs deletion induces transcriptional activation and abnormal translocation of an ectopic S region to compete with upstream S regions

To further address the potential mechanism of the reduction of CSR to various S regions upon the complete deletion the 3’IgH CBEs, we employed GRO-seq to analyze transcription of sequences downstream of the 3’IgH CBEs in WT and complete 3’IgH CBEs-deleted nonstimulated splenic B cells. This analysis revealed that 30kb region just downstream of 3’IgH CBEs was highly activated transcriptionally in non-stimulated, complete 3’IgH CBEs-deleted splenic B cells (Fig. S5A), suggesting that the 3’IgHRR may activate transcription of this downstream IgH region in the absence of the 3’IgH CBEs. Indeed, in contrast to the reduction transcription of various S regions (excluding Sγ1) in αCD40/IL4 or LPS/αIgD-dextran-stimulated splenic B cells, transcription of this immediately downstream IgH region was substantially increased in the absence of the 3’IgH CBEs (Fig. 3 A-D; Fig. S2 and S3). Moreover, our 3C-HTGTS data showed that the iEμ-Sμ locale in the CSRC had greatly increased interactions with this transcriptionally-activated region just downstream of the IgH locus upon deletion of the 3’IgH CBEs (Fig. S4 C-F), indicating that, in the absence of the 3’IgH CBEs, this highly transcribed downstream region participates in substantial synapsis with Sμ within the CSRC (Fig. S4 A, B, D and F).

Transcription across the region downstream of IgH was increased in both sense and antisense directions (Fig. 3 A and C; Fig. S2 and S3), creating convergent transcription known to facilitate AID targeting (25). Notably, αCD40/IL4 treatment of complete 3’IgH CBEs-deleted splenic B cells induced the aberrant translocations across the first 6kb of this 30kb transcribed sequence just downstream of the normal 3’IgH CBEs location, as indicated by junctions between Sμ and sequences across this region that accounted for nearly 1% of all CSR-related junctions (Fig.1 B and C; Fig. S1D; Table S1). In addition, LPS/αIgD-dextran treatment of complete 3’IgH CBEs deleted splenic B cells also induced aberrant translocations between Sμ and sequences within this 6kb sequence downstream of the normal 3’IgH CBEs location that accounted for nearly 3% of all CSR-related junctions (Fig. 2 A and B; Fig. S1E; Table S2). Moreover, more than 90% of these Sμ junctions mapped to known AID target motifs within the 6kb sequence downstream of the normal 3’IgH CBEs location (Fig. 1D; Fig. 2C), consistent with their joining to Sμ via a CSR-related mechanism. The results are also striking, since the level of CSR to the 3’IgH sequences in LPS/αIgD-dextran-treated splenic B cells is similar to that of CSR to the 6kb Sγ2a sequence which has an 1.6-fold higher density of AID target motifs with a much higher percentage of the canonical AGCT motif (Fig. S1F). Thus, these studies indicate that the 3’IgH CBEs prevent the region just downstream of them from becoming transcriptionally activated, synapsing with Sμ in the CSRC, and serving as an ectopically-induced S region (“eS”) for CSR and aberrant chromosomal deletions.

## Discussion

A prior study reported that deletion of the first eight 3’IgH CBEs had little effect on the class switching (17). However. we now demonstrate that complete deletion of all ten 3’IgH CBEs significantly decreases germline transcription and CSR of upstream Sγ3, Sγ2b, Sγ2a and Sε regions, albeit to varying degrees, after stimulation with αCD40/IL4 or LPS/αIgD-dextran. In addition, class-switching to IgA in 3’IgH CBEs-deleted CH12F3 cells was also reduced to about 50% of control levels (15). Taken together, these findings indicate the 3’IgH CBEs variably promote CSR to all upstream S regions except Sγ1, likely by focusing 3’IgHRR region transcriptional enhancing activity on the IgH locus, as opposed to being diverted in part to regions downstream of the IgH locus. In the latter context, it is notable that Sγ1 is not affected by the 3’IgH CBEs deletion, consistent with Sγ1 known to be far less dependent on 3’IgHRR for CSR than other S regions (8, 13, 20, 21). Mechanistically our findings also show that 3’IgH CBEs deletion activates an eS region just downstream of 3’IgH CBEs for aberrant convergent transcription, synapsis with Sμ in the CSRC, and CSR-related deletional joining with the donor Sμ, suggesting that, to some degree, this downstream eS might compete for 3’IgHRR activity with the upstream I promoters in the context of promoting aberrant CSR-related rearrangements in the absence of the 3’IgH CBEs. Overall, our findings implicate the 3’IgH CBEs as an insulator that safeguards the integrity of the normal CSR process by isolating the 3’IgHRR (CSRC) activities within the IgH domain and preventing 3’IgHRR off-target activity outside of the IgH domain that leads to aberrant, CSR-mediated chromosomal deletions.

## Materials and Methods

### Generation of targeted ES cells and chimeric mice

All ten 3’IgH CBEs were deleted by standard gene-targeted deletion approach (26) on both alleles of the WT TC1 mouse ES cells, which were derived from a 129/SV mouse. These homozygous 3’IgH CBEs-deleted ES cells were confirmed by PCR genotyping (Fig. S1 A and B). Subsequently, we deleted both copies of the Aicda (AID) gene from the homozygous 3’IgH CBEs-deleted ES cells via Cas9/gRNA approach. gRNA oligonucleotides used for CRISPR/Cas9 were cloned into the pX330 vector (Addgene plasmid ID 42230) (27). For CSR-HTGTS-seq experiments, the 3’IgH CBEs knockout ES cells were used to generate chimeric mice with totally ES-cell-derived mature B and T lymphocytes via our RAG2-deficient blastocyst complementation (RDBC) (18). For GRO-seq and 3C-HTGTS experiments, the 3’IgH CBEs and AID double knockout ES cells were used to generate chimeric mice with RDBC. Mouse work was performed under protocols approved by the Institutional Animal Care and Use Committee at Boston Children’s Hospital.

### Cell culture

Primary splenic B cells were isolated by a CD43-negative selection kit from chimeric mice and cultured in medium R15 (RPMI1640, 15% FBS, L-glutamate, 1 × penicillin and streptomycin). Primary splenic B cell stimulation was performed with αCD40 (1 μg/ml, eBioscience) plus IL4 (20 ng/ml, PeproTech) or with LPS (25 ng/ml, sigma) plus αIgD-dextran (3 ng/ml) for 96 hrs. CSR-HTGTS-seq was performed in AID-proficient cells stimulated for 96 hours (19). GRO-seq and 3C-HTGTS were performed in AID-deficient cells as previously described (11). Previous studies measured the transcription and interaction of S regions after stimulating the cells for 48 hours (10, 28), while the CSR was usually measured after stimulating the cells for longer times (10, 19). To make a better comparison between CSR, S region transcription, and chromatin interactions we assayed all parameters in splenic B cells stimulated for 96 hours.

### CSR-HTGTS-seq and data analysis

CSR-HTGTS-seq libraries generated with a 5’Sμ bait (11, 19) were prepared from primary splenic B cells stimulated with αCD40/IL4 or LPS/αIgD-dextran for 96 hours. 25 μg gDNA from αCD40/IL4 or LPS/αIgD-dextran-stimulated splenic B cells was sonicated (25s ON and 60s OFF, 2 cycles with low energy input) on Diagenode Bioruptor sonicator. The sonicated DNA fragments were amplified by LAM-PCR with biotinylated 5’Sμ primer. The LAM-PCR products were enriched with streptavidin C1 beads (Thermo Fisher Scientific, #65001) for 4 hours at room temperature. The enriched biotin-labelled LAM-PCR products were ligated with adaptor, followed by nested-PCR with barcode primers and tag-PCR with P5-I5 and P7-I7 primers. 500-1000bp tag-PCR products were purified by separation on 1% TAE gel. CSR-HTGTS-seq libraries were sequenced by paired end 150 bp sequencing on a Next-SeqTM550 (Illumina). More details of the method and analysis have been described (11, 19).

Libraries were processed via our published pipeline (29) and mapped against the AJ851868/mm9 hybrid genome as described previously (30). Data were analyzed and plotted after removing duplicates (11, 19). Each experiment was repeated three times for statistical analyses. The junction numbers within different S regions, as well as the percentage analysis of different S region junctions with respect to total junctions within the C_H_-containing portion of the IgH are listed in the Supplementary Table 1 and 2. Primers used for CSR-HTGTS-seq are listed in Supplementary Table 3.

### 3C-HTGTS

3C-HTGTS analyses (31) were performed on AID^-/-^ mature splenic B cells stimulated with αCD40/IL4 or LPS/αIgD-dextran for 96 hours as previously described (11). 10 million cells were collected and crosslinked with 2% formaldehyde for 10 minutes at room temperature. Then the crosslinked samples were quenched with glycine at a final concentration of 125 mM and lysed in the 3C lysis buffer (50 mM Tris-HCl, pH 7.5, 150 mM NaCl, 5 mM EDTA, 0.5% NP-40, 1% Triton X-100, protease inhibitors). The nuclei were collected and digested with NlaIII enzyme (NEB, R0125) at 37 °C overnight. The digested nuclei samples were ligated with T4 ligase (Promega, M1801) and incubated overnight at 16 °C. The ligated products were treated with Proteinase K (Roche, #03115852001) at 56 °C overnight for decrosslinking and the 3C templates were extracted by phenol/chloroform. The 3C-HTGTS libraries were then sequenced by paired end 150 bp sequencing on Next-Seq™550 (Illumina). More details of the method have been described (11, 31). All the 3C-HTGTS libraries were size-normalized to 370000 total junctions for comparison. For 3C-HTGTS bait interaction frequency analysis, we counted the number of junctions within the indicated bait-interacting locales for both control and experimental groups. For bar graph presentation, the junction number recovered from control groups was normalized to represent 100% and relative experimental values are listed as a percentage of control values (Supplementary Table 1 and 2). Primers used for 3C-HTGTS are listed in Supplementary Table 3. Each experiment was repeated three times for statistical analyses.

### GRO-seq analysis

GRO-seq libraries were prepared from AID^-/-^ mature splenic B cells stimulated with αCD40/IL4 or LPS /αIgD-dextran for 96 hours as described (11). 10 million cells were permeabilized with the fresh made buffer (10 mM Tris-HCl pH 7.4, 300 mM sucrose, 10 mM KCl, 5 mM MgCl2, 1 mM EGTA, 0.05% Tween-20, 0.1% NP40 substitute, 0.5 mM DTT, protease inhibitors and Rnase inhibitor) and resuspended in 100 μl of storage buffer (10 mM Tris-HCl pH 8.0, 25% (V/V) glycerol, 5 mM MgCl2, 0.1 mM EDTA and 5 mM DTT). The nuclear run-on reaction was performed by adding 100 μl of 2X run-on mix (5 mM Tris-HCl PH 8.0, 2.5 mM MgCl2, 0.5 mM DTT, 150 mM KCl, 0.5 mM ATP, 0.5 mM CTP, 0.5 mM GTP, 0.5 mM BrUTP, RNase inhibitor, 1% sarkosyl) at 37 °C for 5 min. RNA was extracted by Trizol. Hydrolysation was performed by adding NaOH at a final concentration of 0.2 N on ice for 18 min, and followed by quenching with ice-cold Tris-HCl pH 6.8 and exchanging buffer bia Bio-Rad P30 columns. Then the purified RNA was incubated with Br-dU antibody-conjugated beads (Santa Cruz biotechnology, sc-32323-ac) for 1 hour. The enriched run-on samples were incubated with RppH (NEB, M0356S) for 1 hour and hydroxyl repair with T4 PNK (NEB, M0201S) for another 1 hour. RT-PCR was performed after the 5’ and 3’ RNA adaptor ligation. The cDNA template was subjected to making GRO-seq libraries by two rounds of PCR. 200-500 bp PCR products from the first round of PCR were purified by separation on 2.5% TAE gel and subjected to the second round of PCR. The final PCR products were further purified by size-selection with SPRIselect beads (Beckman Coulter, B23318). GRO-seq libraries were sequenced via paired end 150 bp sequencing on a Next-Seq™550 and normalized to a coverage of 10 million 100nt reads for display. Transcriptional activity of specific regions was calculated as Reads Per Million Reads (RPM) (Supplementary Table 1 and 2). Each experiment was repeated three times for statistical analyses.

### Statistical analysis

An unpaired two-tailed student t-test was used to examine the significant difference between samples. At least three repeats were done for each statistical analysis. Quantitative data = mean ± s.d., *n.s*. indicates *p*> 0.05, * p≤ 0.05, ** p≤ 0.01, *** p≤ 0.001.

### Data deposition

CSR-HTGTS-seq, GRO-seq and 3C-HTSTS sequencing data analyzed here has been deposited in the GEO database under the accession numbers GSE152193.

## AUTHOR CONTRIBUTIONS

X.Z., and F.W.A. designed the study; X.Z. received the original 3’IgH CBEs-deleted ES cells from H.S.Y. and deleted *Aicda* gene in these ES cells to generate AID-deficient 3’IgH CBEs-deleted ES cells for GRO-seq and 3C-HTGTS studies. A.M.C.-W injected these ES cells for RDBC. X.Z. isolated splenic B cells from chimeric mice for *in vitro* stimulation and performed CSR-HTGTS-seq, GRO-seq and 3C-HTGTS assay; N.K. designed some of the bioinformatics pipelines; X.Z., and F.W.A analyzed and interpreted data; X.Z., and F.W.A. designed figures and wrote the manuscript.

The authors declare no competing financial interests.

## ACKNOWLEDGMENTS

We thank Alt lab members for contribution to the study, particularly Ming Tian and Hwei-Ling Cheng for the advice with ES cell culture, Jianqiao Hu for data uploading. This work was supported by NIH Grant R01AI077595 to F.W.A. F.W.A is an investigator of the Howard Hughes Medical Institute.

## SUPPLEMENTARY MATERIALS

**Figure S1.**
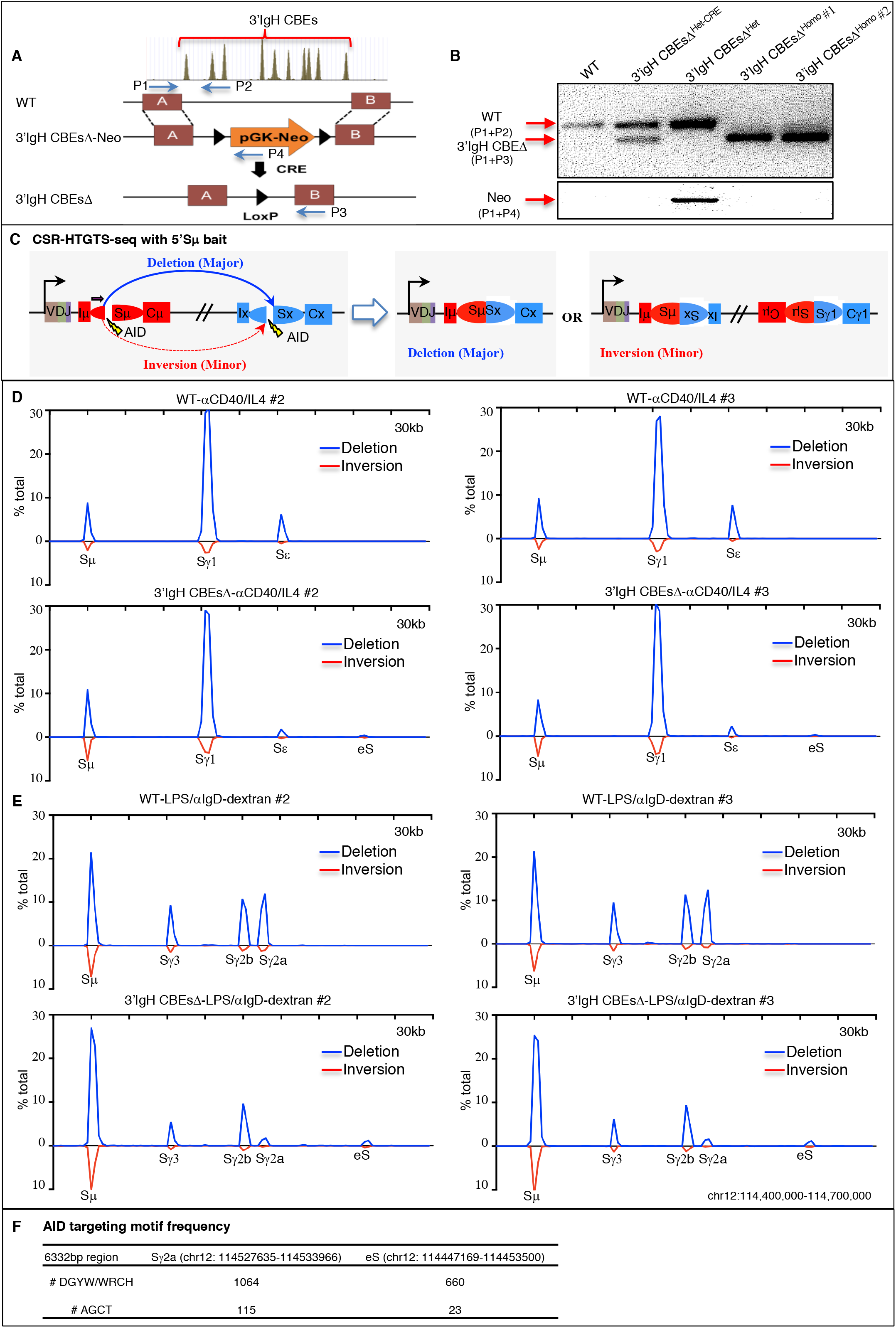
3’IgH CBEs deletion decreases CSR to most upstream S region and induces aberrant translocation to the eS region. (A) Illustration of the targeting strategy used to generate the 3’IgH CBEs-deleted TC1 ES cells. Primer1 (P1) and primer2 (P2) were used for amplifying the WT band. Primer1 (P1) and Primer3 (P3) were used for amplifying the 3’IgH CBEs-deleted band. Primer1 (P1) and primer4 (P4) were used for amplifying the part of pGK-neo band. (B) PCR genotyping to confirm the 3’IgH CBEs-deleted ES clones. (C) Illustration of the detection of CSR by CSR-HTGTS-seq with 5’Sμ bait. As indicated, the vast majority of CSR events are deletional, with an upstream end of an Sμ DSB joining to the downstream end of an acceptor S region DSB. (D) Additional two repeats of CSR-HTGTS-seq data shown in Fig. 1B for αCD40/IL4-stimulated WT and 3’IgH CBEs-deleted splenic B cells. The blue lines indicate deletional joining and the red lines indicate inversional joining. (E) Additional two repeats of CSR-HTGTS-seq data shown in Fig. 2A for LPS/αIgD-dextran-stimulated WT and 3’IgH CBEs-deleted splenic B cells. The blue lines indicate deletional joining and the red lines indicate inversional joining. (F) AID-targeting-motif frequency analysis of 6332bp Sγ2a and eS region.

**Figure S2.**
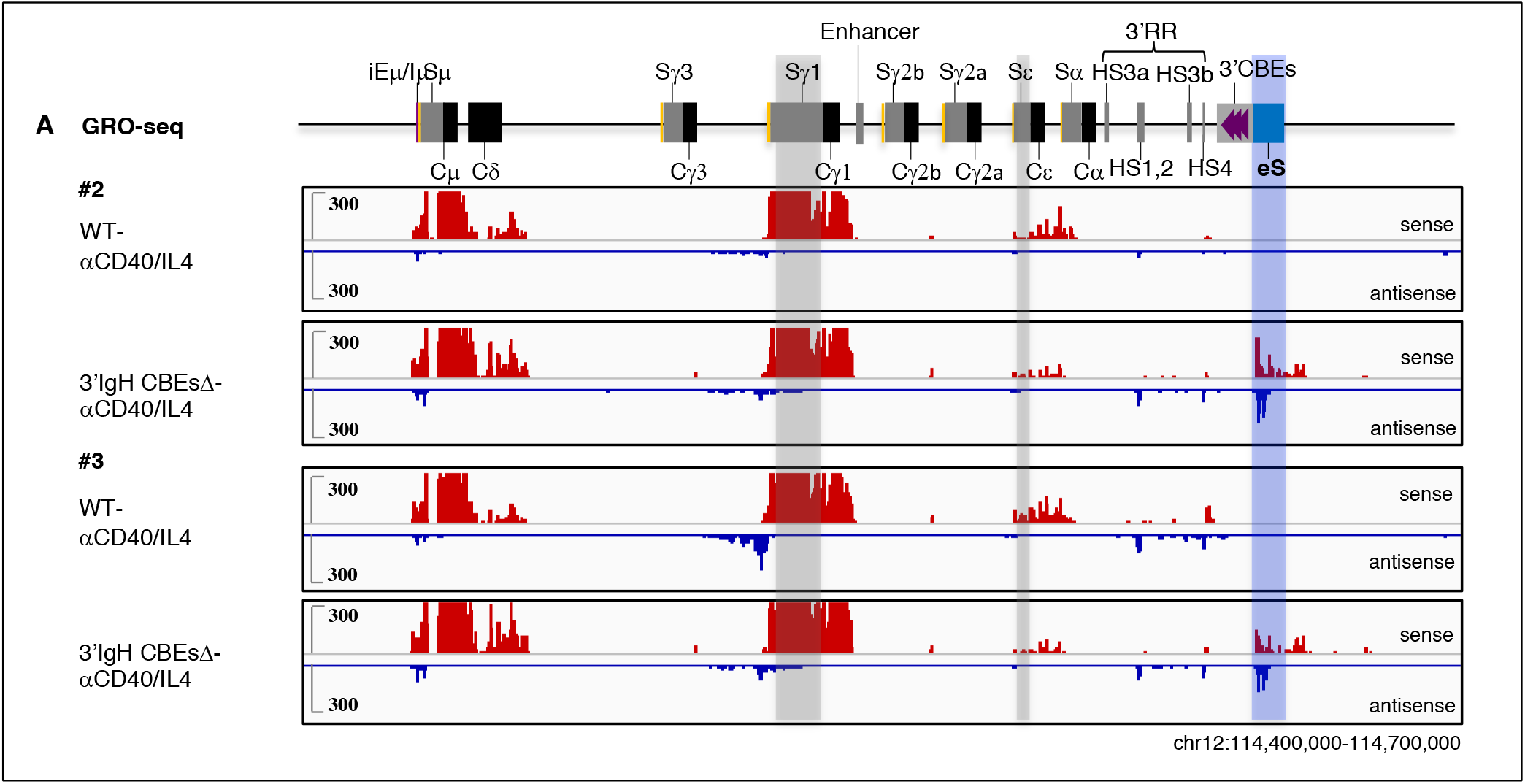
3’IgH CBEs deletion decreases Sε transcription after stimulation with αCD40/IL4 and induces transcription across the downstream eS region. (A) Additional two repeats of GRO-seq data shown in Fig. 3A for αCD40/IL4-stimulated WT and 3’IgH CBEs-deleted splenic B cells. Sense transcription is shown above in red and antisense transcription is shown below in blue lines. Grey bars highlight the Sγ1 and Sε. A blue bar highlight the ectopic S region (labeled as “eS”) just downstream of 3’IgH CBEs.

**Figure S3.**
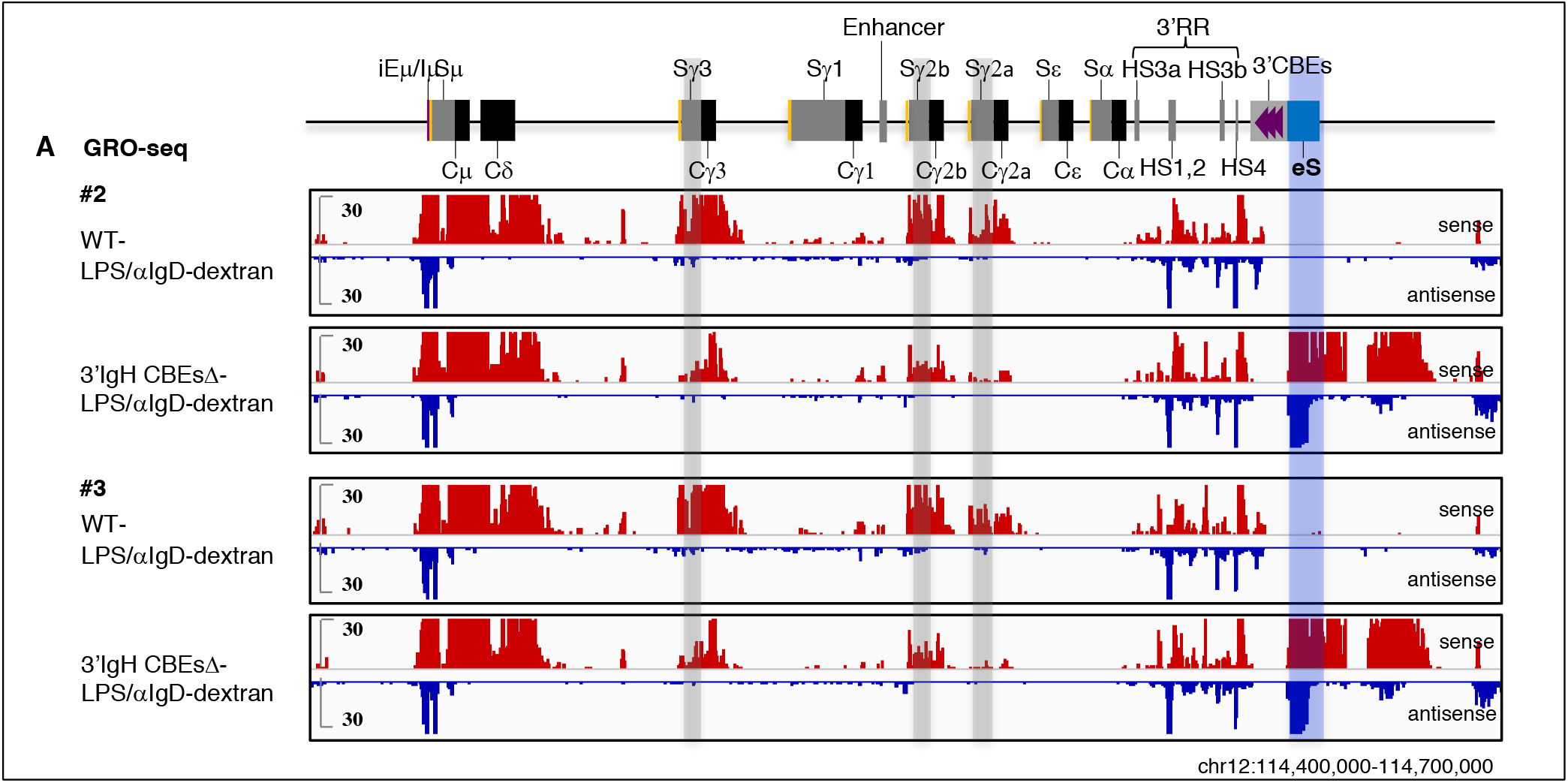
3’IgH CBEs deletion decreases Sγ3, Sγ2b and Sγ2a transcription after stimulation with LPS/αIgD-dextran and induces transcription of the eS region. (A) Additional two repeats of GRO-seq data shown in Fig. 3C for LPS/αIgD-dextran-stimulated WT and 3’IgH CBEs-deleted splenic B cells. Sense transcription is shown above in red and antisense transcription is shown below in blue lines. Grey bars highlight the Sγ3, Sγ2b, Sγ2a, HS3a, HS1,2, HS3b, HS4 and 3’IgH CBEs. A blue bar highlight the ectopic S region (labeled as “eS”) just downstream of 3’IgH CBEs.

**Figure S4.**
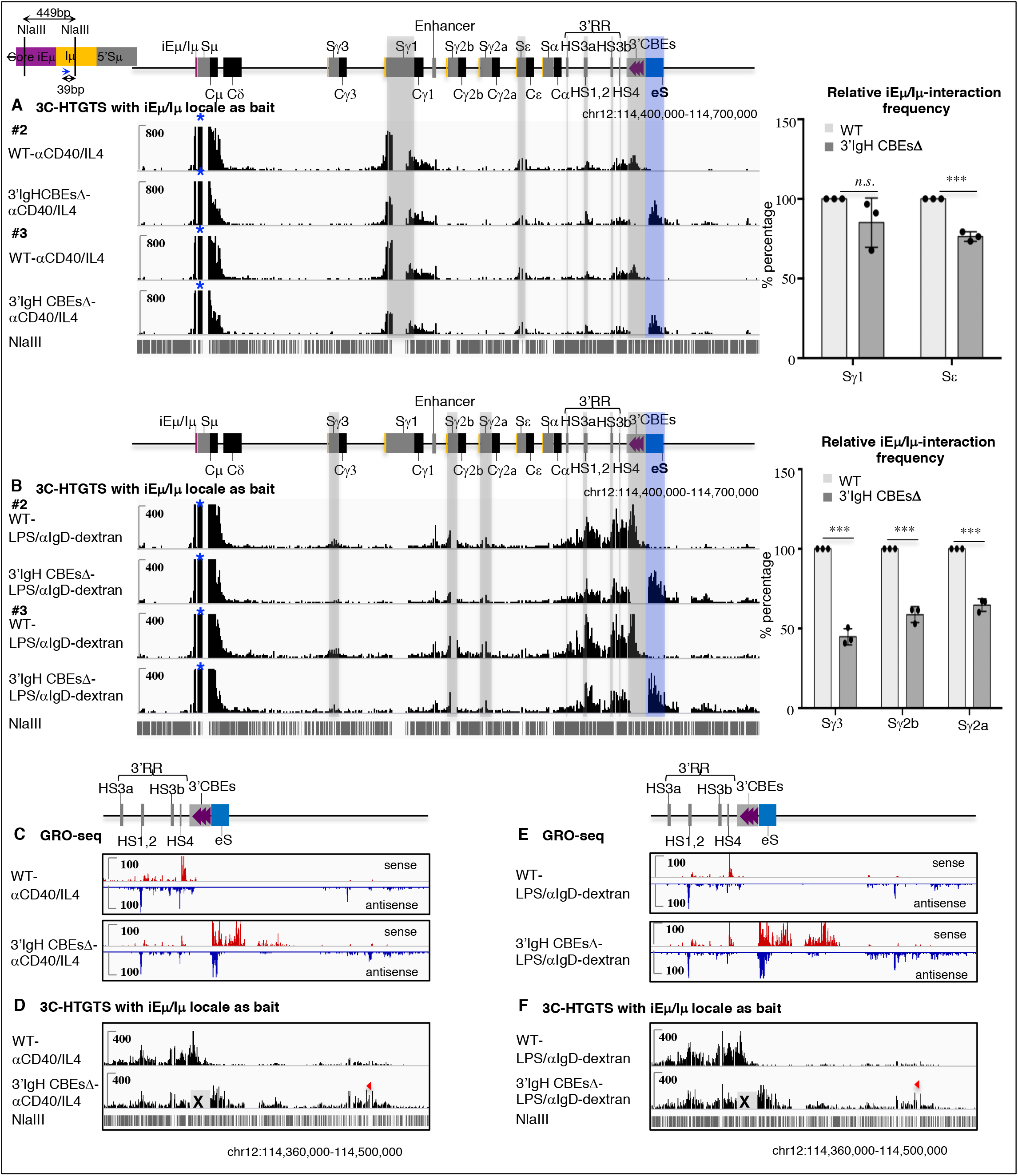
3’IgH CBEs deletion decreases most S-S synapsis and induces Sμ-eS synapsis for abnormal rearrangement. (A) Left: Additional two repeats of 3C-HTGTS data shown in Fig. 4A for αCD40/IL4-stimulated WT and 3’IgH CBEs-deleted splenic B cells. Right: Bar graph shows the relative iEμ-Sμ interaction frequency with Sγ1 and Sε in αCD40/IL4-stimulated splenic B cells. Data represents mean ± s.d. from three independent repeats. *P* values were calculated via unpaired two-tailed *t*-test, *n.s*. indicates*p*> 0.05, *** p≤ 0.001. The raw data for this bar graph is summarized in Table S1. (B) Left: Additional two repeats of 3C-HTGTS data shown in Fig. 4E for LPS/αIgD-dextran-stimulated WT and 3’IgH CBEs-deleted splenic B cells. Right: Bar graph shows the relative iEμ-Sμ interaction frequency with Sγ3, Sγ2b and Sγ2a in LPS/αIgD-dextran-stimulated splenic B cells. Data represents mean ± s.d. from three independent repeats. *P* values were calculated via unpaired two-tailed *t*-test, *** p≤ 0.001. The raw data for this bar graph is summarized in Table S2. (C and E) Magnified GRO-seq profiles to better reveal the transcriptional activation of the region downstream of 3’IgH CBEs after 3’IgH CBEs deletion. (D and F) Magnified 3C-HTGTS profiles to better reveal the interaction between iEμ/Iμ bait with the impediments downstream of 3’IgH CBEs after 3’IgH CBEs deletion.

**Figure S5.**
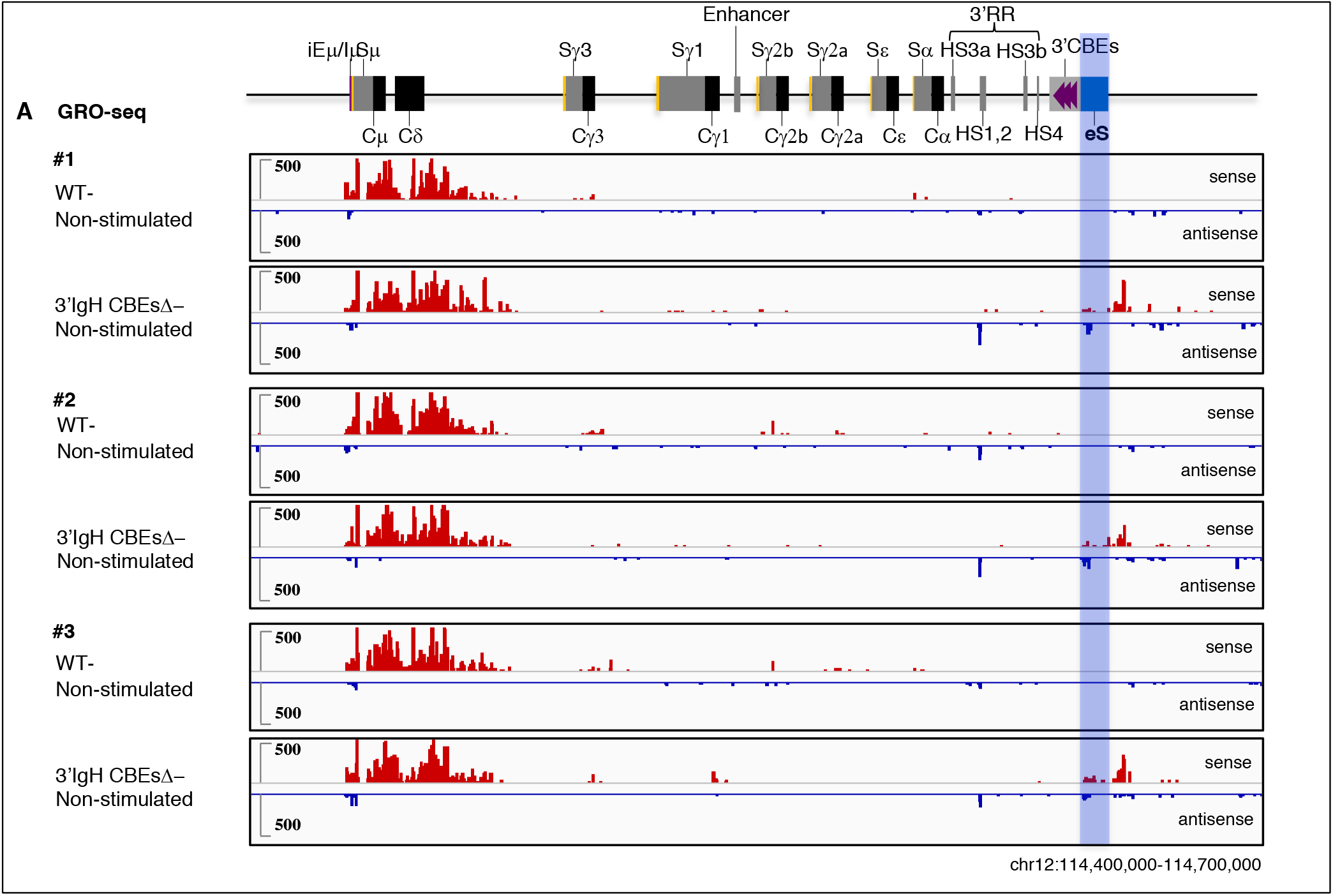
3’IgH CBEs deletion induces transcription of the eS region in non-stimulated splenic B cells. A) GRO-seq profiles of IgH locus from non-stimulated WT and 3’IgH CBEs-deleted splenic B cells. Sense transcription is shown above in red and antisense transcription is shown below in blue lines. A blue bar highlight the ectopic S region (labeled as “eS”) just downstream of 3’IgH CBEs.

**Table S1.** The effect of 3’IgH CBEs deletion on switching, transcription and interaction of different S regions in WT and 3’IgH CBEsΔ splenic B cells stimulated with αCD40/IL4.

| αCD40/IL4 | IgH | S<sub>γ1</sub> | S<sub>ε</sub> | eS |
| --- | --- | --- | --- | --- |
| WT #1 | 9090 (100%) | 6722 (73.9%) | 966 (10.6%) | 0 (0%) |
| WT #2 | 11284 (100%) | 8660 (76.8%) | 1023 (9.1%) | 0 (0%) |
| WT #3 | 8915 (100%) | 6498 (72.9%) | 1022 (11.5%) | 0 (0%) |
| 3'IgH CBEsΔ #1 | 5077 (100%) | 3785 (74.6%) | 185 (3.6%) | 35 (0.7%) |
| 3'IgH CBEsΔ #2 | 6320 (100%) | 4767 (75.4%) | 183 (1.9%) | 54 (0.9%) |
| 3'IgH CBEsΔ #3 | 3750 (100%) | 2920 (77.9%) | 119 (3.2%) | 27 (0.7%) |

| αCD40/IL4 | S<sub>γ1</sub> | S<sub>ε</sub> | eS |
| --- | --- | --- | --- |
| WT #1 | 1109 | 128 | 0 |
| WT #2 | 1558 | 168 | 0 |
| WT #3 | 1593 | 175 | 0 |
| 3'IgH CBEsΔ #1 | 1184 | 71 | 105 |
| 3'IgH CBEsΔ #2 | 1080 | 65 | 75 |
| 3'IgH CBEsΔ #3 | 1103 | 63 | 76 |

| αCD40/IL4 | S<sub>γ1</sub> | S<sub>ε</sub> | eS |
| --- | --- | --- | --- |
| WT #1 | 100 | 100 | 100 |
| WT #2 | 100 | 100 | 100 |
| WT #3 | 100 | 100 | 100 |
| 3'IgH CBEsΔ #1 | 91 | 79 | 1226 |
| 3'IgH CBEsΔ #2 | 97 | 73 | 1230 |
| 3'IgH CBEsΔ #3 | 68 | 76 | 2075 |

**Table S2.** The effect of 3’IgH CBEs deletion on switching, transcription and interaction of different S regions in WT and 3’IgH CBEsΔ splenic B cells stimulated with LPS/αIgD-dextran.

Relative utilization of different S regions in WT and 3'IgH CBEsΔ splenic B cells stimulated with αCD40/IL4.
| LPS/algD-dextran | IgH | S<sub>γ</sub>3 | S<sub>γ</sub>2b | S<sub>γ</sub>2a | eS |
| --- | --- | --- | --- | --- | --- |
| WT #1 | 3933 (100%) | 564 (14.3%) | 857 (21.8%) | 902 (22.9%) | 1 (0.02%) |
| WT #2 | 4940 (100%) | 680 (13.7%) | 1039 (21.0%) | 1138 (23.0%) | 0 (0%) |
| WT #3 | 6097 (100%) | 875 (14.4%) | 1285 (21.1%) | 1385 (22.7%) | 0 (0%) |
| 3'IgH CBEsΔ #1 | 2087 (100%) | 175 (8.4%) | 338 (16.2%) | 71 (3.4%) | 57 (2.7%) |
| 3'IgH CBEsΔ #2 | 2836 (100%) | 216(7.6%) | 471 (16.6%) | 101 (3.6%) | 77 (2.7%) |
| 3'IgH CBEsΔ #3 | 2958 (100%) | 247 (8.4%) | 479 (16.2%) | 95 (3.2%) | 69 (2.4%) |

| LPS/algD-dextran | S<sub>γ</sub>3 | S<sub>γ</sub>2b | S<sub>γ</sub>2a | eS |
| --- | --- | --- | --- | --- |
| WT #1 | 69 | 24 | 11 | 0 |
| WT #2 | 40 | 16 | 6 | 0 |
| WT #3 | 46 | 16 | 6 | 0 |
| 3'IgH CBEsΔ #1 | 7 | 5 | 1 | 65 |
| 3'IgH CBEsΔ #2 | 7 | 6 | 1 | 61 |
| 3'IgH CBEsΔ #3 | 8 | 7 | 1 | 69 |

| LPS/algD-dextran | S<sub>γ</sub>3 | S<sub>γ</sub>2b | S<sub>γ</sub>2a | eS |
| --- | --- | --- | --- | --- |
| WT #1 | 100 | 100 | 100 | 100 |
| WT #2 | 100 | 100 | 100 | 100 |
| WT #3 | 100 | 100 | 100 | 100 |
| 3'IgH CBEsΔ #1 | 41 | 61 | 60 | 1809 |
| 3'IgH CBEsΔ #2 | 51 | 62 | 67 | 1643 |
| 3'IgH CBEsΔ #3 | 43 | 53 | 67 | 2288 |

**Table S3.**
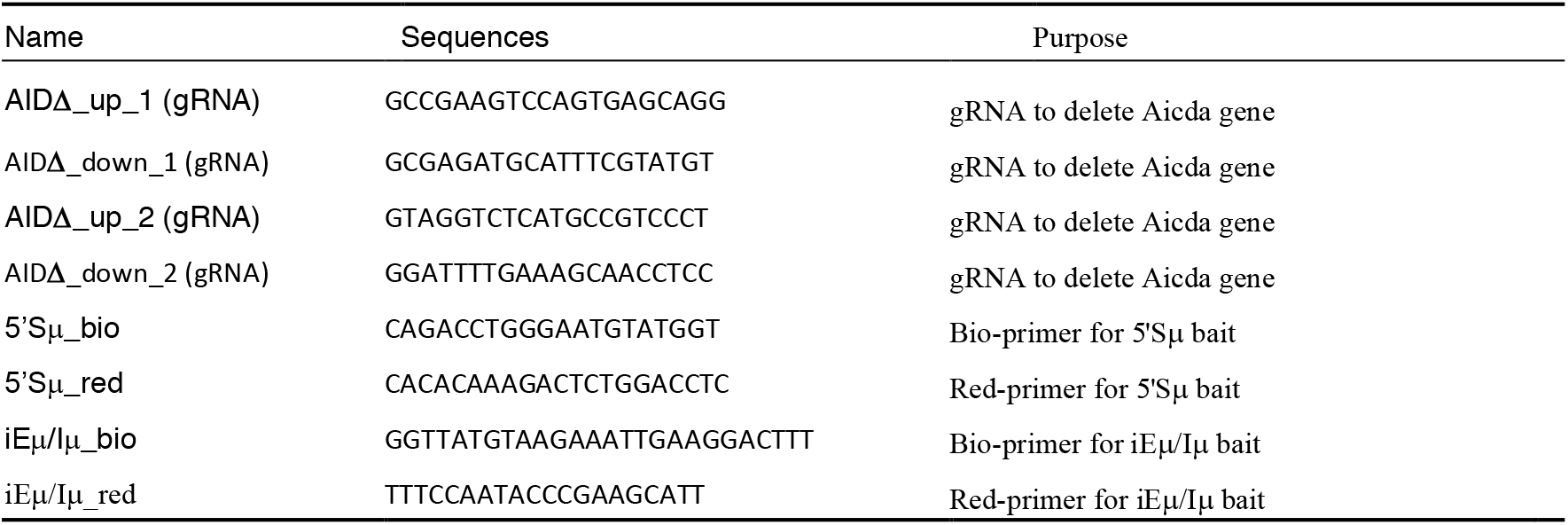
List of oligos.

